# LLM-powered Functional Gene Set Summarization with genesetGPT

**DOI:** 10.64898/2026.07.27.741117

**Authors:** Jack R. Leary, Samantha Pattey, Rhonda Bacher

## Abstract

Transcriptomics datasets generated using next-generation sequencing techniques such as single cell RNA-sequencing (scRNA-seq) and spatially-resolved transcriptomics (SRT) allow researchers to study patterns in gene expression across celltypes, temporal processes, and spatial organization at ever-higher resolutions and depths. scRNA-seq analyses produce gene expression profiles and celltype-specific gene sets that require annotation to provide biological meaning, a process that has traditionally relied on the manual interpretations of clinical scientists. Similarly, SRT experiments typically require subjective, time-consuming annotation of spatial domains. Recent advances in large language model (LLM) methods offer opportunities to assist in the interpretation of such datasets. Many current LLM-based approaches aim to annotate transcriptomics-derived gene sets by integrating information from publicly available and online biological resources. While these approaches can be effective, they often struggle when presented with weakly-related or fully uncorrelated genes, sometimes inferring and justifying biological relationships that are not supported by existing literature. Additionally, the quality of LLM-generated interpretations is dependent on the provision of appropriate biological context and careful prompt design, both of which can present significant barriers to effective use. To address these limitations we propose genesetGPT, an efficient, LLM-based framework that emphasizes both curated biological context and iterative prompt construction, thus enabling realistic summarization of heterogeneous gene sets at scale. genesetGPT is implemented as an open-source Python package available for download at https://github.com/jr-leary7/genesetGPT.

## 1. Introduction

One of the primary outputs of transcriptomic data analysis are gene sets, which arise across a variety of contexts, including differential expression (DE) analysis between celltypes or experimental conditions, identification of spatially variable genes (SVG), or inference of transcription factor activity signatures. In cases where gene sets are small or composed of well-characterized genes, their biological interpretation is relatively straightforward. However, modern transcriptomics studies using high resolution technologies such as single-cell RNA-sequencing (scRNA-seq) and spatially-resolved transcriptomics (SRT) capture multiple, overlapping axes of biological variation, yielding gene sets ranging in size from hundreds to thousands of genes that are not readily interpretable. To address this challenge, methods that summarize gene sets in terms of functional pathways or biological processes are commonly used. For example, gene set enrichment analysis (GSEA), one of the foundational methods for functional interpretation of gene sets, uses permutation testing to assess whether a given gene set is significantly enriched in predefined biological pathways or processes[1]. While GSEA and related methods have proven broadly useful, they share a fundamental limitation in that gene function is highly context-dependent, varying across tissues, cell types, and cell states. For example, the role of gene *APOE*, which broadly functions throughout the human body as a modulator of lipid transportation, differs substantially between the brain [2] and peripheral blood [3].

Despite the development of numerous alternatives to classical GSEA, many popular enrichment methods available in both R [4–6] and Python [7, 8] do not enable user provision of fine-grained contextual information such as celltype composition, system state, or the preprocessing and analysis steps employed when generating the gene set. As a result, enrichment results can be overly vague or even wholly irrelevant to the underlying biology [9]. One context-aware R package, clusterProfiler [10], implements AI-based interpretation guided by user-provided biological background knowledge, though its results are crucially based on previously-identified enriched pathways instead of richer information describing the function(s) of each individual gene.

More recently, natural language processing (NLP) methodologies have been applied to text mining problems by mapping textual datasets to numerical vector spaces using both general embedding frameworks such as Word2Vec [11], and domain-specific ones like BioWordVec [12]. While such embedding representations allowed for downstream statistical learning tasks, early models often suffered from limitations related to data sparsity, polysemy, and out-of-vocabulary generalization [13]. These issues lead to the introduction of the transformer architecture [14] and large language models (LLMs), which have since scaled significantly in both parameter volumes and training dataset sizes [15] while empirically demonstrating capabilities of completing wide varieties of complex reasoning tasks [16].

These recent advances in the summarization of large textual corpora have led researchers to develop LLM-based methods for gene set summarization [17–19], which have shown some promise in describing gene set functions [20]. TALISMAN [17] implements a command line interface (CLI) based on OpenAI’s GPT models in order to generate putative gene set names and summaries, though it only queries one of two functional databases - either RefSeq [21] or gene ontology (GO) descriptions [22]. IAN [19] uses Google’s Gemini [23] to analyze differentially expressed (DE) gene sets. However, IAN’s summarizations rely solely on *a priori* aggregated enrichment analysis results from tools like Enrichr [24] or clusterProfiler [10], and not on literature-validated information concerning each individual gene’s function(s). In general, current LLM-based gene set summarization methods are hampered by either reliance on a single model provider, generation of gene set summaries using incomplete reference data, or lack of a simple programmatic interfaces.

Beyond extant tools’ inputs and outputs, several more general difficulties persist. First, LLM results are inherently stochastic, even when relevant hyperparameters are fixed [25, 26]. This variability can be exacerbated when the LLM is not provided with adequate biological context. Second, constructing effective, parsimonious prompts is often nontrivial [27]. Each prompt must be precisely engineered and formatted in order to ensure that the model understands the biological question clearly and is given enough background information to generate a useful response, without being provided extraneous details.

To address the linked issues of growing ‘omics analysis output sizes and lack of result interpretability we propose genesetGPT, an LLM-based method for guided, structured, functional summarization of gene sets derived from modern transcriptomics datasets (**Fig. 1**). The open-source genesetGPT Python package is freely available for download via GitHub at https://github.com/jr-leary7/genesetGPT.

**Figure 1.**
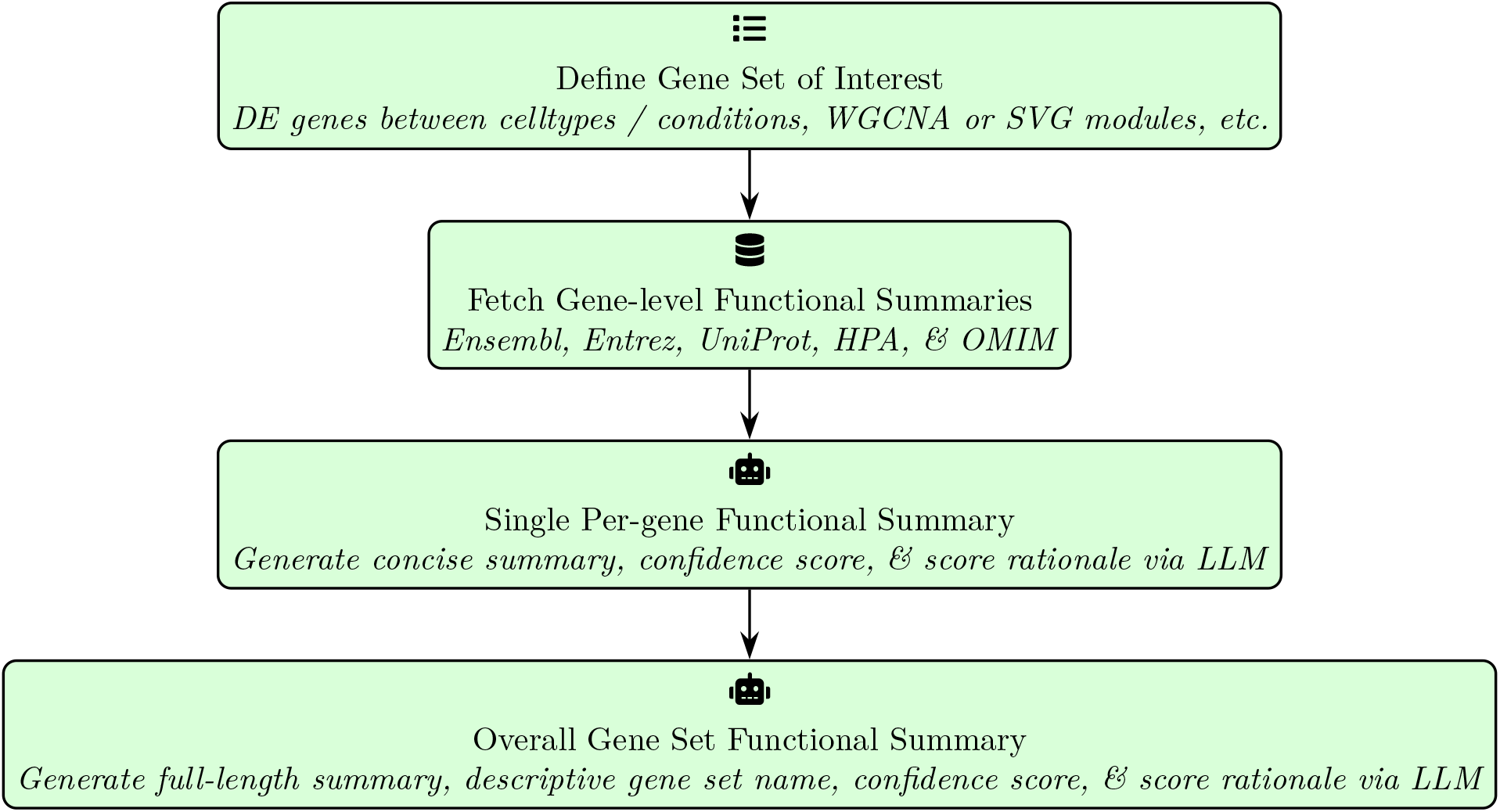
Central steps of functional gene set inference using genesetGPT. Both the per-gene LLM summaries and the overall gene set-level LLM summary may be used to guide further downstream analyses and inform biological interpretations.

## 2. Methods

### 2.1 Building a gene function database

In order to precisely guide the backend LLM towards generating a realistic, literature-based summary of each individual gene and then each gene module, we first fetch summaries of each gene’s function from multiple different databases. We begin by pulling the Ensembl and Entrez IDs associated with each HGNC symbol in the gene set, along with the corresponding gene description, biotype, and chromosomal location from the Ensembl database [28]. Next, we query the Online Mendelian Inheritance of Man (OMIM) [29] database’s mapping of MIM IDs to the other aforementioned gene IDs, and use that information to identify any phenotypes associated with variations in each gene. We then pull the NCBI Entrez summary [30] for each gene, after which we leverage the Human Protein Atlas to determine which diseases and GO:BP terms each gene is involved in. Finally, we fetch data from UniProtconcerning each gene’s canonical protein product along with any corresponding functional summaries.

#### 2.2 Prompt construction

After collecting all the data sources for each gene in the user’s gene set of interest, genesetGPT uses a rule-based system to collate each piece into a single, per-gene prompt to be passed to the backend LLM in order to create a concise summary of each gene’s function.

genesetGPT’s prompting routine is broken down into two distinct parts: the user prompt and the system prompt. The user prompt contains the textual information that is passed to the LLM during each individual API call i.e., what data the end user would like the LLM to synthesize, whereas the system prompt specifies an overarching, concrete role for the LLM, optional constraints on the format of its responses, and any biological context relevant to the dataset being analyzed and the hypotheses being investigated [33] (**Box 1**).

In addition to the user and system prompts, we include an additional prompt instructing the model to generate a unique, per-annotation confidence score ranging from 0-1 that estimates the reliability of each summary, along with a brief, 2-4 sentences in length, rationale justifying the score.

When summarizing entire gene modules, we also prompt the LLM to determine a concise and distinctive name for the module based on its overall functional summary and confidence in its annotation. This module name provides researchers with a concrete starting point for drilling down further into their data and examining the LLM’s conclusions.

With respect to computational efficiency, genesetGPT is trivially scalable to larger gene sets. Since the per-gene prompt construction and LLM summarization routines are I/O-bound as opposed to CPU-bound, we are able to leverage asynchronous execution of the underlying API calls to parallelize both operations via shared-memory multithreading [35, 36]. This allows the number of workers employed to execute the underlying API calls in parallel to be much greater than the number of logical cores on the end user’s machine, with the primary constraint on overall execution speed being the rate at which those functions are able to return results.

**Box 1.**

An example gene module summarization system prompt. The prompt provides a specific role for the model, defines explicit output constraints, and offers biological context concerning how the data were generated and processed. When passing this information to the backend LLM’s Python API, Markdown-style syntax is used in order to both prevent LLM confusion as to which sections of the system prompt are relevant to which end user directives, and refine LLM outputs by aligning with the immense amount of Markdown-formatted text in the model’s train set [34].

\# Role

You are an experienced computational biologist with extensive knowledge of transcriptomics analyses such as single-cell RNA-seq and spatially-resolved transcriptomics. When generating responses, you consider the statistical, computational, and biological angles of the question at hand. Your responses are precise and detailed without being too overly technical.

\# Strict Constraints

In all responses, do not utilize any means of referring to a gene other than its HGNC symbol e.g., never use ‘Neurogranin’ to refer to the gene NRGN. Do not reference any information that is not explicitly present in the user prompt. Lastly, do not hesitate to express and quantify your uncertainty if the module’s genes have highly diverse or unclear functionalities.

\# Biological Context

The system being studied is the healthy human cerebral cortex, and the data were sequenced using 10X Genomics Visium V1. Quality control, preprocessing, and downstream analyses were performed with a standard workflow based on the Scanpy and Squidpy Python packages. Lastly, the genes in this module have been sorted in decreasing order of spatial significance via the Moran’s I test i.e., genes whose expression is more statistically significantly dependent on spatial location appear earlier in the user prompt’s bulleted list and vice versa.

### 2.3 Dataset processing

#### 2.3.1 Human cerebral cortex

After downloading the Visium V1 human cortex dataset from 10X Genomics [37], we performed basic spot- and gene-level quality control (QC) by removing spots with a sequencing depth of less than 1000 and genes expressed in fewer than five spots. Next, a set of 3000 HVGs was identified using the “seurat_v3” method implemented in the Scanpy Python package [38]. The raw counts were then depth-normalized, log1p-transformed, and scaled. Initial linear dimension reduction was performed using PCA [39, 40] after subsetting the normalized, scaled counts matrix to include only the HVGs. A KNN graph with ĸ = 20 neighbors was then estimated using the top 30 PCs as input, after which the Leiden algorithm [41] with a resolution of *r* = 0.5 was used to sort the graph into clusters. The top 30 PCs were further dimension-reduced to 2D for visualization purposes via UMAP [42]. Next, a spatial neighborhood graph with ĸ = 6 neighbors was generated prior to determining a ranking of the HVGs by their spatial variability as estimated by the Moran’s I statistic [43]. The top 1000 genes having Benjamini-Hochberg [44] adjusted *p*-values lower than 0.05 were classified as SVGs and used to further subset the normalized counts matrix, which was then transposed such that genes were treated as rows and spots as columns. This matrix was scaled and reduced to 30 dimensions via PCA, after which a gene-level KNN graph was estimated using the Euclidean distance andĸ = 20 neighbors. This graph was then partitioned into discrete clusters via the Leiden algorithm with resolution *r* = 0.0025, resulting in a total of four distinct SVG modules. Each module was then scored per-spot based on the normalized counts using the `sc.tl.score_genes()` Scanpy function. Finally, genesetGPT summarization was performed per-gene and per-module using Anthropic’s Claude Haiku v4.5 and Opus v4.5 models, respectively. All of the genesetGPT functions were run with otherwise default settings, with the exception of slight increases of the value of the `n_max_tokens` parameter to 1000 for the gene-level summarization, and 6000 for the module-level summarization.

#### 2.3.2 Artificial null gene module experiment

In order to interrogate the fidelity of the gene module summaries generated by genesetGPT, we designed an empirical test of our method’s ability to confidently determine whether or not a gene module exhibits biologically-plausible shared functionalities. We utilized the human cortex SRT dataset with its four “true” SVG modules estimated as described previously, to which we added a fifth artificial null SVG module created by randomly sampling 250 genes from the set of all non-SVGs. We then compared the LLM-generated module summary, estimated confidence score, and confidence score rationale assigned to the artificial gene module with those of the true SVG modules.

Next, we repeated the summarization process 50 times for each module and recorded the genesetGPT-generated confidence scores at each iteration. We then estimated the mean, standard deviation, and range of the scores for each module in order to quantify their stability.

#### 2.3.3 Human bone marrow hematopoesis

After downloading the *CD34*+ human bone marrow hematopoesis dataset [45], we leveraged the CellRank2 Python package [46] to identify potential cell fate driver genes. First, after removing cells annotated as common lymphoid progenitors (CLPs), we derived a kernel-based, cell-cell transition probability matrix based on the *a priori* estimated Palantir pseudotime [45]. Next, we used the GPCCA method [47] to compute and manually define one initial (Hematopoetic Stem Cell) and four terminal (Erythrocyte, Megakaryocyte, Monocyte, and DC) cell macrostates based on both the computational results and known biology. We then estimated every cell’s probability of differentiating into each of the four possible cell fates based on both their relative locations in the 2D t-SNE [48] embedding of the trajectory and their pseudotime estimates. We next employed the permutation test implemented in CellRank2’s `compute_lineage_drivers()`function to estimate the significance of the Pearson correlations [49] between each cell’s four fate probability estimates and each gene’s normalized expression in those cells in order to identify putative driver genes. The relevant `n_perms` parameter was set to 1000 for each cell fate’s correlation testing routine, and results were filtered to only include genes with positive correlations having Benjamini-Hochberg [44] adjusted *p*-values less than 0.01. Finally, we utilized a default genesetGPT workflow with the Claude Haiku v4.5 (gene-level) and Opus v4.5 (module-level) backends to summarize the top 50 lineage-driving genes for each cell fate.

## Results

### 3.1 Distinct SVG modules in the human cerebral cortex

The human brain is a complex, strictly spatially-structured organ, making high-throughput SRT an excellent tool for characterizing its underlying biological mechanisms. An immediate side effect of said complexity, however, is that it can be quite difficult to describe precisely how different genes interact with one another functionally in their spatial contexts. This issue can be exacerbated when assaying tissues with spot-level methods like 10X Genomics Visium that aggregate gene expression counts across small groups of cells, versus newer technologies that quantify spatial gene expression within each cell. As such, the analysis of SVG modules derived from a healthy human cerebral cortex sample assayed using 10X’s Visium V1 technology was determined to be a suitable test of genesetGPT’s module summarization capabilities.

After preprocessing the data, we obtained five distinct clusters of spots that approximately represented cortical layers and showed clear separation in gene expression space (**Fig. 2A-B**). The set of 1000 SVGs exhibited complex and strongly spatially-dependent patterns of gene expression (**Fig. 2C**). For example, the second most spatially-significant gene, *MOBP*, has been canonically characterized in multiple model organisms as intrinsically involved in myelination [50, 51], and is thus also differentially expressed in oligodendrocytes [52]. After partitioning the SVG set into four discrete modules, *MOBP* was assigned to SVG Module 1, which was composed of 190 unique genes (**Fig. 2D**). However, a module composed of even 190 genes - whether they are *a priori* annotated or not - is realistically too large for a human researcher to functionally synthesize in a reasonable amount of time with a high degree of accuracy. Ergo, we applied genesetGPT to each of the SVG modules in order to accelerate the process of spatial niche annotation.

**Figure 2.**
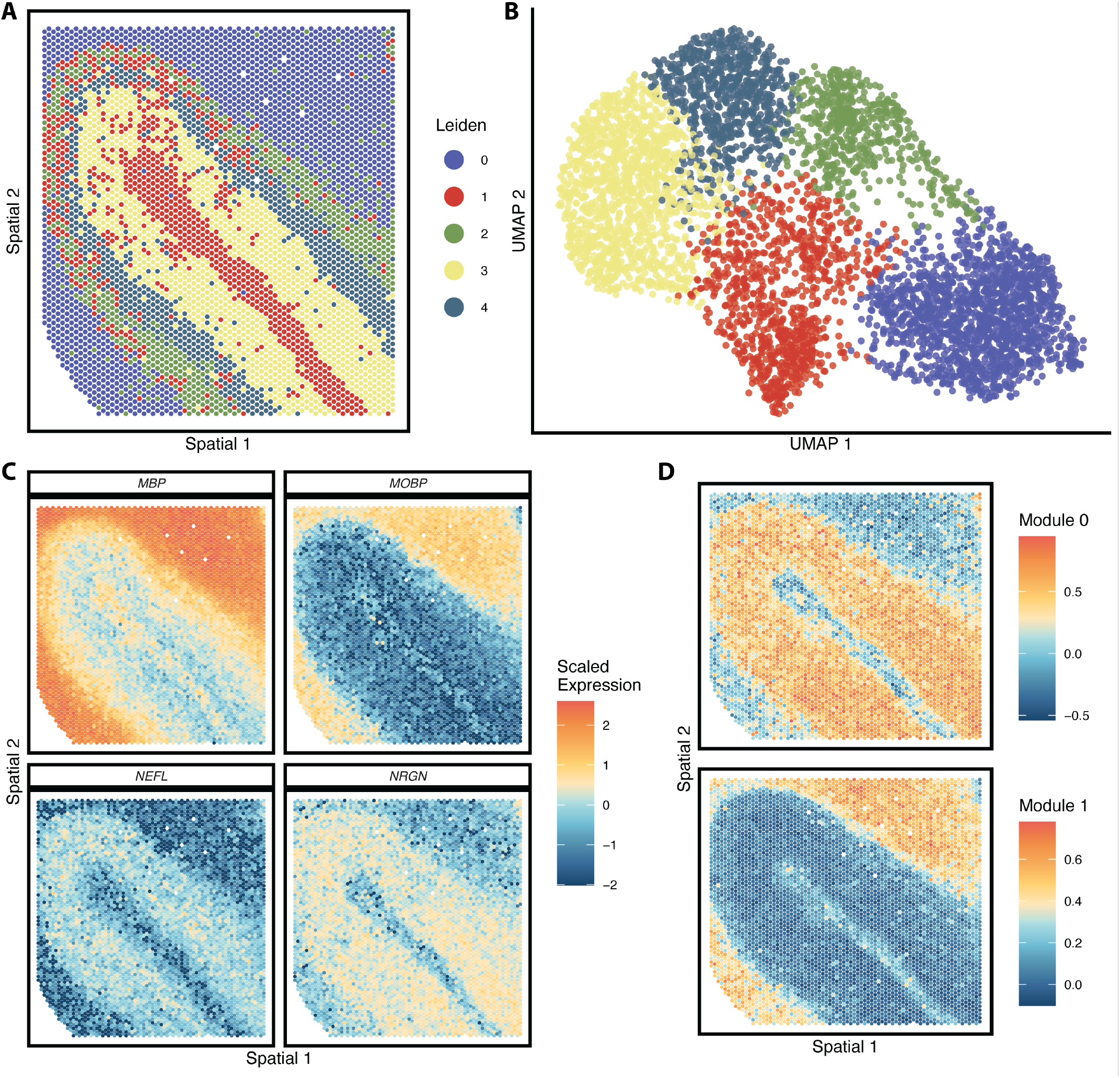
Clustering of SVGs in the human cerebral cortex dataset into spatial modules prior to summarization with genesetGPT. **(A)** Spatial scatterplot of 4901 spots colored by Leiden cluster IDs derived from the 1000 SVG expression profiles. **(B)** UMAP embedding scatterplot with spots colored by SVG expression-based cluster IDs as in **(A). (C)** Spatial scatterplots colored by the scaled, normalized gene expression of the top four SVGs. **(D)** Spatial scatterplots of estimated per-spot module scores for SVG Module 0 & Module 1.

After performing gene- and gene module-level summarization with genesetGPT we obtained the following description for SVG Module 1, which the LLM named “Myelination and White Matter Program” (**Box 2**). The backend LLM’s accurate characterization of *MOBP* as being relevant to cortical myelination was followed by an explicit reference to the module’s additional involvement in oligodendrocyte differentiation. genesetGPT’s precise identification of such myelination-related SVGs as *MOBP*, functionally similar genes like *MOG* [53], and the oligodendrocyte maturation-regulating transcription factor *MYRF* [54] indicate that genesetGPT’s internal data acquisition and subsequent structured prompt construction resulted in a nuanced and realistic summarization of the module as a whole.

The LLM-estimated confidence score for SVG Module 1 was 0.78, and the corresponding concise rationale proposed by the LLM for that score was grounded in the underlying biology without being overconfident or hyperbolic in its conclusions (**Box 3**).

**Box 2.**

Functional summary of SVG Module 1 in the healthy human cortex SRT dataset generated by genesetGPT using Anthropic’s Claude Haiku v4.5 as the backend LLM.

This 190-gene module represents a comprehensive program of myelination, myelin maintenance, and associated white matter biology in the healthy human cortex. Core myelin structural proteins (*MOG, MBP* via *MYRF* regulation, *MAG, CNP, ERMN, MOBP, OPALIN, PLLP, UGT8, FA2H, CLDN11*) are co-expressed with oligodendrocyte differentiation factors (*NKX6-2, MYRF, BCAS1*) and lipid metabolism enzymes essential for myelin biogenesis (*ABCA8, ENPP2, PLA2G4D*). The module integrates vascular and immune surveillance components (*HLA-DPA1, HLA-DRA, HLA-DRB1, HLA-E, CCL2, CCL4, CD44, TLR2, LACC1*), consistent with the metabolic and immune demands of mature white matter, alongside neuronal support functions including synaptic regulation (*BDNF, NPY, HTR3A, PENK*), ion homeostasis (*AQP1, KCNH8, PIEZO2*), and axonal integrity (*CNTN2, MAG*). Spatially heterogeneous expression of these genes likely reflects distinct white matter tracts, cortical layers with varying myelination density, and regional differences in oligodendrocyte populations and their metabolic/immune states within the cortex.

**Box 3.**

genesetGPT’s rationale justifying its estimated confidence score of 0.78 that accompanied its summarization of SVG Module 1’s biological functionality.

The annotation is strongly supported by the prevalence of well-characterized myelin structural proteins and oligodendrocyte differentiation factors with consistent functional descriptions across authoritative databases. However, the inclusion of 190 genes introduces considerable functional heterogeneity, with substantial subsets devoted to vascular function, immune regulation, and other processes whose specific roles in the cortical white matter context remain partially uncertain. The confidence score reflects high confidence in the dominant myelination theme but accounts for incomplete characterization of some genes and contextual ambiguities in interpreting spatial variability in adult cortex tissue.

### 3.2 genesetGPT exercises caution on a false gene set

When summarizing and scoring the artificial null SVG module generated using QC-passing non-SVGs detected in the human cortex SRT dataset, genesetGPT convincingly demonstrated its ability to identify and quantify uncertainty without hallucinating functional relationships. Compared to the lowest estimated confidence score among the four true SVG modules (0.78), genesetGPT scored the artificial module 76.28% lower (**Table 1**). In addition, the paragraph-length summary generated by genesetGPT for the artificial null SVG module included explicit language specifying that the gene set was unlikely to be a cohesive gene program with a detectable purpose (**Box 4**). In particular, genesetGPT identified a “lack of clear pathway enrichment” among the 250 genes, and concluded by positing that the artificial module could just be the result of “technical noise”. Lastly, the LLM assigned the name “Heterogeneous Cellular Maintenance Functions”, the vagueness of which was consistent with the randomly selected nature of the artificial null module.

**Table 1.** Estimated confidence scores for the artificial null set experiment gene modules, where Module 4 is composed of the 250 randomly selected non-SVGs.

| Module | Assigned Name | Confidence Score |
| --- | --- | --- |
| 0 | Cortical Neuronal Function Module | 0.925 |
| 1 | White Matter Myelination and Neuroimmune | 0.78 |
| 2 | Cortical Synaptic Transmission Module | 0.88 |
| 3 | Cortical Synaptic Signaling Network | 0.875 |
| 4 | Heterogeneous Cellular Maintenance Functions | 0.185 |

**Box 4.**

The genesetGPT summary for the artificial null SVG module.

This gene set comprises a highly heterogeneous collection of 250 genes with diverse molecular functions spanning transcriptional regulation, DNA repair, mitochondrial metabolism, immune signaling, and cellular housekeeping processes. A notable subset includes numerous zinc finger proteins (*ZNF14, ZNF215, ZNF219, ZNF227, ZNF350-AS1, ZNF432, ZNF454, ZNF630, ZNF682, ZNF689, ZNF761, ZNF784, ZNF823*) that function as transcriptional regulators, alongside chromatin-modifying factors such as *HMGN5, BRMS1L*, and *SAP30BP*. Several genes participate in protein quality control and degradation pathways, including ERAD components (*DERL1*), ubiquitin ligase machinery (*KBTBD4, RNF166, FBXL8*), and ribosome quality control (*ANKZF1*). Mitochondrial function is represented by genes involved in oxidative phosphorylation (*NDUFB3*), mitochondrial translation (*MRPL17, MRPL48, TFB1M*), and protein import (*TIMM50*). The set also contains genes involved in cell cycle regulation (*CDC20, SPC25, CDKN2A, PSRC1*) and DNA repair (*XPC, BRIP1, NEIL2, EME1*). The functional diversity and lack of clear pathway enrichment suggest this module may represent baseline cellular maintenance processes or technical noise rather than a coherent biological program.

The estimated confidence for the artificial null SVG module was 0.185, where genesetGPT’s score justification noted that many genes in the set lacked concrete functional annotations and the indicated pathways were too disjoint to be biologically meaningful (**Box 5**). Finally, genesetGPT again raised the possibility of the module being the result of technical noise.

The genesetGPT-generated confidence scores for each true SVG module exhibited low variability, thus indicating a high degree of reliability (**Table 2**). The confidence scores for the artificial null module (Module 4) had the lowest mean (0.164) and widest range (0.2), accurately reflecting the random, unconnected nature of the genes that comprised the set.

**Table 2.**
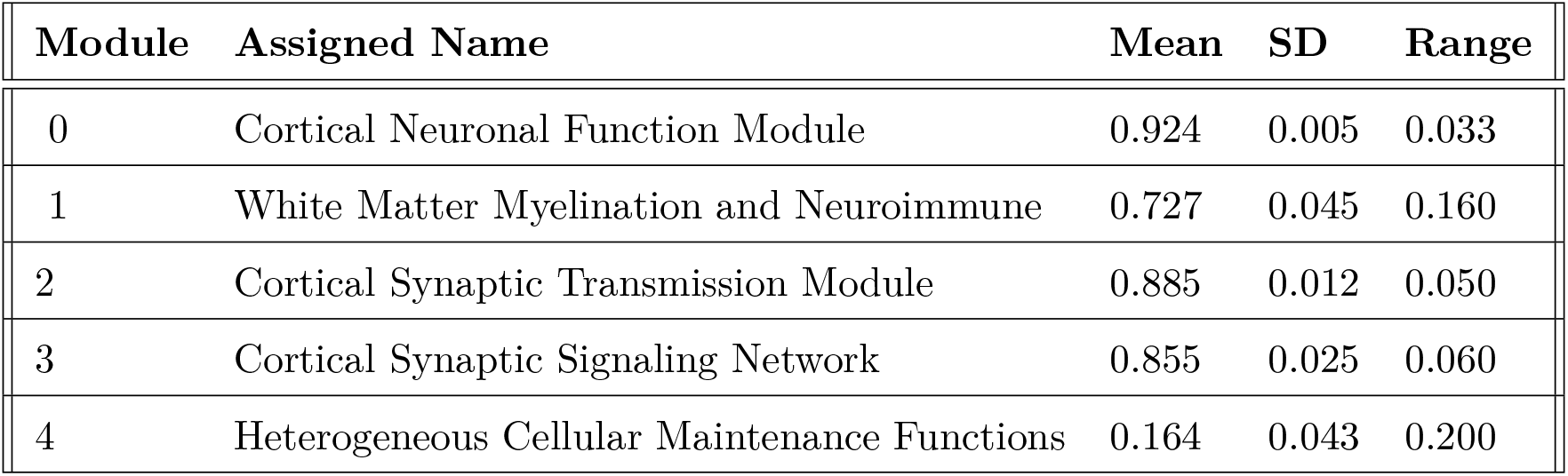
Summary statistics measuring the variability of the cerebral cortex dataset’s SVG module confidence scores estimated using genesetGPT.

| Module | Assigned Name | Mean | SD | Range |
| --- | --- | --- | --- | --- |
| 0 | Cortical Neuronal Function Module | 0.924 | 0.005 | 0.033 |
| 1 | White Matter Myelination and Neuroimmune | 0.727 | 0.045 | 0.160 |
| 2 | Cortical Synaptic Transmission Module | 0.885 | 0.012 | 0.050 |
| 3 | Cortical Synaptic Signaling Network | 0.855 | 0.025 | 0.060 |
| 4 | Heterogeneous Cellular Maintenance Functions | 0.164 | 0.043 | 0.200 |

In terms of computational efficiency, fetching each of the gene annotations from the online databases then building each of the unique, per-gene user prompts took 12.25 minutes with the `n_workers` parameter set to 20. The gene-level LLM summary generation and confidence scoring routine completed in 12.45 minutes using the same settings.

**Box 5.**

The genesetGPT justification of its estimated confidence score for the artificial null SVG module.

The extremely low confidence reflects the profound functional heterogeneity within this 250-gene set, which lacks a dominant unifying biological theme. The genes span disparate processes including transcription, DNA repair, mitochondrial function, immune signaling, and lipid metabolism without clear pathway enrichment. Many genes are poorly characterized or represent tissue-inappropriate expression for cortical tissue, suggesting this module may capture technical artifacts or non-specific housekeeping functions rather than a biologically meaningful program.

The sequentially-executed overall LLM summarization of the five SVG modules ran in 2.25 minutes. Each analysis step was run on a 2020 Intel i5 MacBook Pro with 16GB of memory.

### 3.3 Megakaryocyte lineage-driving genes during hematopoesis

We next characterized a scRNA-seq-derived set of genes, each of which was statistically significantly correlated with commitment to the Megakaryocyte cell fate during hematopoesis in human *CD34*+ bone marrow cells [45, 46] (**Fig. 3A-B**). After preprocessing the scRNA-seq counts, we identified putative driver genes for each of the four broad terminal cell states and applied genesetGPT to distinguish lineage-specific drivers of commitment to the Megakaryocyte cell fate from those of the closely-linked Erythrocyte cell fate.

**Figure 3.**
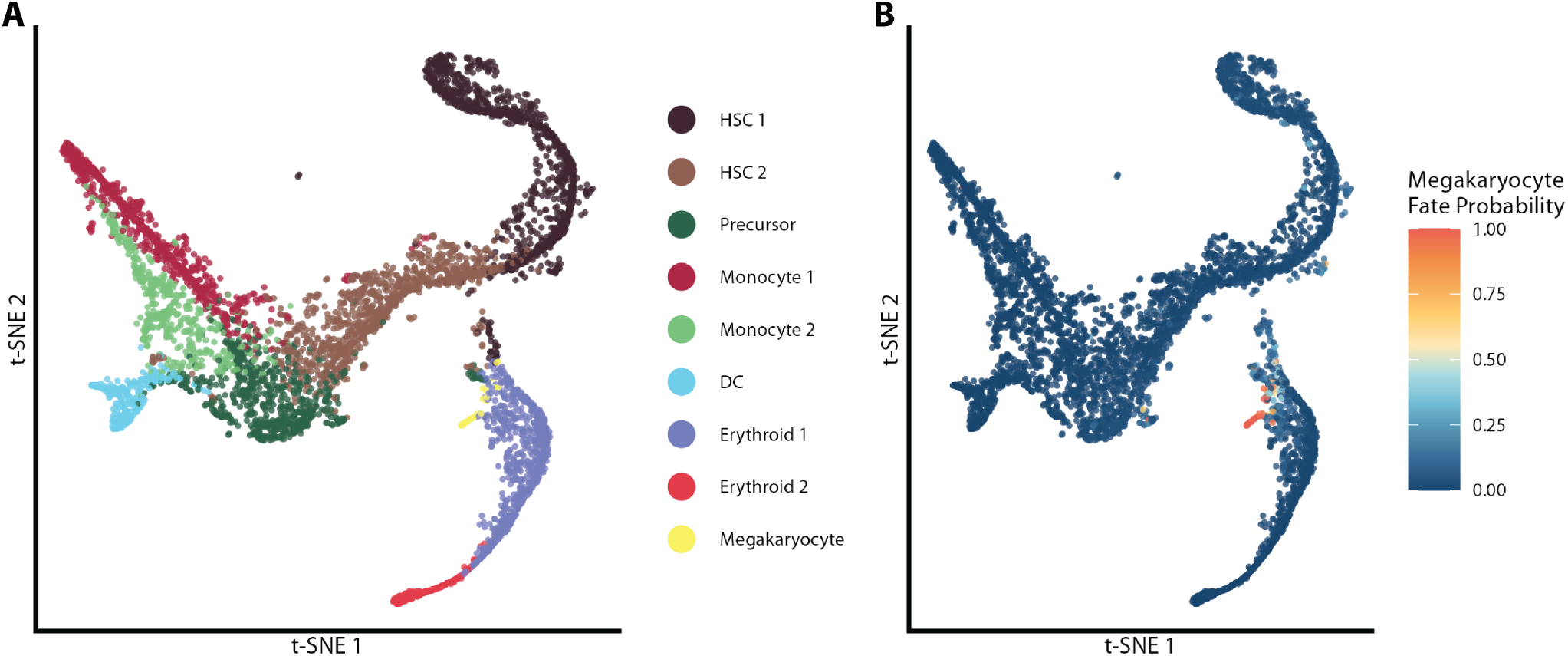
Megakaryocyte cell fate specification during human bone marrow hematopoesis. **(A)** t-SNE embedding of *CD34*+ bone marrow cells colored by their annotated celltypes. **(B)** t-SNE embedding as in (**A**) colored by each cell’s estimated probability of commitment to the Megakaryocyte cell fate

genesetGPT’s summary for the Megakaryocyte cell fate’s lineage drivers highlighted canonical Megakaryocyte-specific genes such as *PF4* [55], and *PPBP* [56]. The assigned module name was “Megakaryocyte-Adhesion-Signaling Module”, accurately reflecting the gene module’s statistical association with Megakaryocyte fate commitment (**Box 6**).

The estimated confidence score that accompanied genesetGPT’s functional summary was 0.52, and the accompanying rationale indicated clear consistency with known Megakaryocyte biology as well as measured hesitation towards unilaterally assigning a single role to the module (**Box 7**).

Efficiency-wise, genesetGPT fetched gene summaries and constructed user prompts for the top 50 driver genes of each of the four potential cell fates in 4.9 minutes with the `n_workers` parameter set to 20, after which performing the per-gene LLM summarization took only 44 seconds with the same value of `n_workers`. As described previously, each function call was executed on a 2020 Intel i5 MacBook Pro with 16GB of memory.

## 4 Discussion

genesetGPT allows researchers to rapidly and accurately summarize the functions of computationally-derived gene modules by precisely constructing biologically-guided per-gene and per-module LLM prompts. Each underlying prompt is based solely on current biological information queried from comprehensive databases such as Ensembl, UniProt, and the Human Protein Atlas. Additionally, genesetGPT provides estimated confidence scores and accompanying score justifications for every individual annotation, thus allowing researchers to easily determine the degrees of functional coherence exhibited by their gene modules.

Our first case study involved the analysis of SVG modules derived from a healthy human cortex SRT dataset. Primarily, we identified an SVG module that genesetGPT characterized as being defined by genes strongly related to both myelination and oligodendrocyte differentiation. genesetGPT’s overall summary was backed up by references to literature-validated genes such as *MOBP* and *MOG*. Additionally, genesetGPT provided a succinct, measured rationale for its confidence in its own summarization without hallucinating a singular, overarching function for the SVG module by pinpointing distinct sources of uncertainty in its final annotation.

We next tested genesetGPT on an artificial null module composed of genes that were randomly sampled from the set of non-SVGs in the human cerebral cortex. genesetGPT was able to efficiently identify and describe the clear lack of functional cohesion in the artificial null module, assigning and justifying a low confidence score. We also quantified the variability of the genesetGPT-estimated module confidence scores, showing that the true SVG modules exhibited scores with high means and low variances, and vice versa for the artificial null module.

**Box 6.**

genesetGPT’s functional summary of the top 50 putative driver genes that were significantly associated with commitment to the Megakaryocyte cell fate.

This 50-gene module represents a functionally heterogeneous set with a core theme of megakaryocyte commitment and platelet biology, complemented by broader roles in hematopoietic cell adhesion, calcium signaling, and cytoskeletal dynamics. Multiple genes encode platelet-associated surface receptors and coagulation factors (*GP1BA, GP9, ITGA2B, VWF, SELP, CLEC1B, PF4, PPBP*), while others regulate calcium homeostasis (*SLC24A3, TRPC6, ITPKA*) and vesicular trafficking (*ASAP2, DNM3, STXBP5*) processes essential for differentiation. Cytoskeletal regulators (*ARHGAP6, PDLIM1, TPM1, PLEK*) and transcriptional regulators (*PBX1, NFIB, MED12L*) support lineage specification across multiple fates, while adhesion molecules (*CDH6, CD9, THBS1*) and immune signaling adaptors (*LAT, RGS18, RGS6*) facilitate cell fate determination. However, the module contains substantial functional noise, including genes with primarily neuronal functions (*APBA2, BDNF, DOK6*), musculoskeletal genes (*TNNT1, COL2A1*), sensory genes (*PIEZO2, PKHD1L1*), and metabolic/endocrine genes (*GMPR, SPX, CMTM5, LINC00504*) whose relevance to hematopoietic commitment remains unclear or potentially artifactual.

**Box 7.**

The genesetGPT-generated rationale for its estimated confidence score accompanying the Megakaryocyte fate-specification module’s overall functional summary.

Confidence is moderate-to-low due to significant functional heterogeneity within the 50 genes. While a coherent subset (approximately 15-20 genes) strongly supports megakaryocyte commitment and platelet biology with high specificity, roughly one-third of the module comprises genes with well-established non-hematopoietic functions (neuronal, musculoskeletal, sensory) whose presence suggests either genuine context-dependent biology or potential false positives from the computational pipeline. The inclusion of poorly characterized genes (*LINC00504, CMTM5, PLXDC2*) and metabolic genes (*GMPR, SPX*) further reduces interpretive confidence.

Our final case study concerned the analysis of a set of putative cell fate-specifying genes for Megakaryocytes in a human bone marrow hematopoesis scRNA-seq dataset. genesetGPT was able to succinctly characterize the Megakaryocyte lineage, even when provided with only its top 50 commitment-driving genes. genesetGPT also miti-gated overconfidence by readily identifying individual genes whose functions appeared unrelated to Megakaryocyte development, despite the fact that the Megakaryocyte driver gene module was constructed based on a very small set of gene expression profiles (comprising just 1.2% of the 5292 total cells).

A limitation of the current genesetGPT implementation is that it only supports the summarization of gene sets derived from *H. sapiens* samples. However, adding support for datasets generated from other well-studied model organisms, particularly the ubiquitous *M. musculus*, would prove valuable to a large cohort of researchers. Additionally, while genesetGPT currently fully supports Anthropic’s Claude and OpenAI’s GPT APIs as LLM backends for structured gene set summarization, multiple other rapidly-improving LLM platforms such as Google’s Gemini [23] and Meta’s Llama [57] are widely-used. Enabling researchers to choose from an even larger set of backend LLMs would improve genesetGPT’s reproducibility.

Lastly, there remains work to be done pertaining to the estimation of gene- and module-level confidence scores. While we showed that the scores estimated by genesetGPT are empirically reliable and low variance, recent literature has identified potential issues with current confidence-scoring approaches. For example, cheaper, less-complex models are more likely to systematically generate higher confidence scores [58]. This can be partially ameliorated by more precise, biologically-specific prompting of the backend LLM [59], analogous to what we implemented in the genesetGPT Python package. In addition, genesetGPT’s provision of succinct, concrete justifications linked to it’s estimated confidence scores allows researchers to easily gauge whether the gene- and module-level summaries are biologically plausible with respect to the underlying system being studied.

## 5 Acknowledgments

## 5.1 Data & code availability

The genesetGPT Python package is available to download as free, open source software via GitHub at https://github.com/jr-leary7/genesetGPT. All analyses performed for this manuscript are available at https://github.com/jr-leary7/genesetgpt-analyses. Each processed dataset used in our case study analyses is available for download in AnnData format through Zenodo at https://doi.org/10.5281/zenodo.21307654.

## 5.2 Author contributions

J.R.L designed the real data analyses and implemented the software package with assistance from S.P. S.P. performed the artificial null gene module experiment. J.R.L. and S.P. drafted the manuscript. R.B. edited the manuscript and guided revisions.

## References

[1] Aravind Subramanian, Pablo Tamayo, Vamsi K. Mootha, et al. “Gene set enrichment analysis: A knowledge-based approach for interpreting genome-wide expression profiles”. Proceedings of the National Academy of Sciences 102.43 (Oct. 2005), pp. 15545–15550.

[2] Abdel Ali Belaidi, Ashley I. Bush, and Scott Ayton. “Apolipoprotein E in Alzheimer’s disease: molecular insights and therapeutic opportunities”. Molecular Neurodegeneration 20.1 (Apr. 2025), p. 47.

[3] Ana B. Martínez-Martínez, Elena Torres-Perez, Nicholas Devanney, et al. “Beyond the CNS: The many peripheral roles of APOE”. Neurobiology of Disease 138 (May 2020), p. 104809.

[4] Liis Kolberg, Uku Raudvere, Ivan Kuzmin, et al. “g:Profiler—interoperable web service for functional enrichment analysis and gene identifier mapping (2023 update)”. Nucleic Acids Research 51.W1 (July 2023), W207–W212.

[5] Zhuorui Xie, Allison Bailey, Maxim V. Kuleshov, et al. “Gene Set Knowledge Discovery with Enrichr”. Current Protocols 1.3 (Mar. 2021), e90.

[6] Gennady Korotkevich, Vladimir Sukhov, Nikolay Budin, et al. Fast gene set enrichment analysis. June 2016.

[7] Alexander Lachmann, Zhuorui Xie, and Avi Ma’ayan. “blitzGSEA: efficient computation of gene set enrichment analysis through gamma distribution approximation”. Bioinformatics 38.8 (Apr. 2022). Ed. by Valentina Boeva, pp. 2356–2357.

[8] Zhuoqing Fang, Xinyuan Liu, and Gary Peltz. “GSEApy: a comprehensive package for performing gene set enrichment analysis in Python”. Bioinformatics 39.1 (Jan. 2023). Ed. by Zhiyong Lu, btac757.

[9] Farhad Maleki, Katie Ovens, Daniel J. Hogan, et al. “Gene Set Analysis: Challenges, Opportunities, and Future Research”. Frontiers in Genetics 11 (June 2020), p. 654.

[10] Shuangbin Xu, Erqiang Hu, Yantong Cai, et al. “Using clusterProfiler to characterize multiomics data”. Nature Protocols 19.11 (Nov. 2024), pp. 3292–3320.

[11] Tomas Mikolov, Kai Chen, Greg Corrado, et al. Efficient Estimation of Word Representations in Vector Space. 2013.

[12] Yijia Zhang, Qingyu Chen, Zhihao Yang, et al. “BioWordVec, improving biomedical word embeddings with subword information and MeSH”. Scientific Data 6.1 (May 2019), p. 52.

[13] Jose Camacho-Collados and Mohammad Taher Pilehvar. “From Word To Sense Embeddings: A Survey on Vector Representations of Meaning”. Journal of Artificial Intelligence Research 63 (Dec. 2018), pp. 743–788.

[14] Ashish Vaswani, Noam Shazeer, Niki Parmar, et al. Attention Is All You Need. 2017.

[15] Jared Kaplan, Sam McCandlish, Tom Henighan, et al. Scaling Laws for Neural Language Models. 2020.

[16] Avinash Patil and Aryan Jadon. Advancing Reasoning in Large Language Models: Promising Methods and Approaches. Version Number: 2. 2025.

[17] Marcin P. Joachimiak, J. Harry Caufield, Nomi L. Harris, et al. Gene Set Summarization using Large Language Models. 2023.

[18] Zhizheng Wang, Qiao Jin, Chih-Hsuan Wei, et al. “GeneAgent: self-verification language agent for gene-set analysis using domain databases”. Nature Methods 22.8 (Aug. 2025), pp. 1677–1685.

[19] Vijayaraj Nagarajan, Reiko Horai, Guangpu Shi, et al. “IAN, an intelligent system for omics data analysis and discovery”. Cell Reports Methods (June 2026), p. 101503.

[20] Mengzhou Hu, Sahar Alkhairy, Ingoo Lee, et al. “Evaluation of large language models for discovery of gene set function”. Nature Methods 22.1 (Jan. 2025), pp. 82–91.

[21] Tamara Goldfarb, Vamsi K Kodali, Shashikant Pujar, et al. “NCBI RefSeq: reference sequence standards through 25 years of curation and annotation”. Nucleic Acids Research 53.D1 (Jan. 2025), pp. D243–D257.

[22] The Alliance of Genome Resources Consortium, Suzanne A Aleksander, Anna V Anagnostopoulos, et al. “Updates to the Alliance of Genome Resources central infrastructure”. GENETICS 227.1 (May 2024). Ed. by V Wood, iyae049.

[23] Gemini Team, Rohan Anil, Sebastian Borgeaud, et al. Gemini: A Family of Highly Capable Multimodal Models. 2023.

[24] Edward Y Chen, Christopher M Tan, Yan Kou, et al. “Enrichr: interactive and collaborative HTML5 gene list enrichment analysis tool”. BMC Bioinformatics 14.1 (Dec. 2013), p. 128.

[25] Berk Atil, Sarp Aykent, Alexa Chittams, et al. Non-Determinism of “Deterministic” LLM Settings. 2024.

[26] Shuyin Ouyang, Jie M. Zhang, Mark Harman, et al. “An Empirical Study of the Non-Determinism of ChatGPT in Code Generation”. ACM Transactions on Software Engineering and Methodology 34.2 (Feb. 2025), pp. 1–28.

[27] Banghao Chen, Zhaofeng Zhang, Nicolas Langrené, et al. “Unleashing the potential of prompt engineering for large language models”. Patterns 6.6 (June 2025), p. 101260.

[28] Sarah C Dyer, Olanrewaju Austine-Orimoloye, Andrey G Azov, et al. “Ensembl 2025”. Nucleic Acids Research 53.D1 (Jan. 2025), pp. D948–D957.

[29] A. Hamosh. “Online Mendelian Inheritance in Man (OMIM), a knowledgebase of human genes and genetic disorders”. Nucleic Acids Research 33.Database issue (Dec. 2004), pp. D514–D517.

[30] Eric W Sayers, Jeffrey Beck, Evan E Bolton, et al. “Database resources of the National Center for Biotechnology Information in 2025”. Nucleic Acids Research 53.D1 (Jan. 2025), pp. D20–D29.

[31] Mathias Uhlén, Linn Fagerberg, Björn M. Hallström, et al. “Tissue-based map of the human proteome”. Science 347.6220 (Jan. 2015), p. 1260419.

[32] The UniProt Consortium, Alex Bateman, Maria-Jesus Martin, et al. “UniProt: the Universal Protein Knowledgebase in 2025”. Nucleic Acids Research 53.D1 (Jan. 2025), pp. D609–D617.

[33] Anna Neumann, Elisabeth Kirsten, Muhammad Bilal Zafar, et al. “Position is Power: System Prompts as a Mechanism of Bias in Large Language Models (LLMs)”. Proceedings of the 2025 ACM Conference on Fairness, Accountability, and Transparency. Athens Greece: ACM, June 2025, pp. 573–598.

[34] Jia He, Mukund Rungta, David Koleczek, et al. Does Prompt Formatting Have Any Impact on LLM Performance? Version Number: 1. 2024.

[35] Robert H. Halstead. “MULTILISP: a language for concurrent symbolic computation”. ACM Transactions on Programming Languages and Systems 7.4 (Oct. 1985), pp. 501–538.

[36] Brian Quinlan. PEP 3148 – futures-execute computations asynchronously. PEP 3148. Python Software Foundation, 2009.

[37] 10X Genomics. Adult Human Brain 1 (Cerebral Cortex, Unknown orientation). June 2020.

[38] F. Alexander Wolf, Philipp Angerer, and Fabian J. Theis. “SCANPY: large-scale single-cell gene expression data analysis”. Genome Biology 19.1 (Dec. 2018), p. 15.

[39] Karl Pearson. “On lines and planes of closest fit to systems of points in space”. The London, Edinburgh, and Dublin Philosophical Magazine and Journal of Science 2.11 (Nov. 1901), pp. 559–572.

[40] H. Hotelling. “Analysis of a complex of statistical variables into principal components.” Journal of Educational Psychology 24.6 (Sept. 1933), pp. 417–441.

[41] V. A. Traag, L. Waltman, and N. J. van Eck. “From Louvain to Leiden: guaranteeing well-connected communities”. Sci Rep 9.1 (Mar. 2019), p. 5233.

[42] Leland McInnes, John Healy, and James Melville. “UMAP: Uniform Manifold Approximation and Projection for Dimension Reduction” (Sept. 2020). Version Number: 3.

[43] P. A. P. Moran. “Notes on Continuous Stochastic Phenomena”. Biometrika 37.1/2 (June 1950), p. 17.

[44] Yoav Benjamini and Yosef Hochberg. “Controlling the False Discovery Rate: A Practical and Powerful Approach to Multiple Testing”. Journal of the Royal Statistical Society Series B: Statistical Methodology 57.1 (Jan. 1995), pp. 289–300.

[45] Manu Setty, Vaidotas Kiseliovas, Jacob Levine, et al. “Characterization of cell fate probabilities in single-cell data with Palantir”. Nature Biotechnology 37.4 (Apr. 2019), pp. 451–460.

[46] Philipp Weiler, Marius Lange, Michal Klein, et al. “CellRank 2: unified fate mapping in multiview single-cell data”. Nature Methods 21.7 (July 2024), pp. 1196–1205.

[47] Bernhard Reuter, Marcus Weber, Konstantin Fackeldey, et al. “Generalized Markov State Modeling Method for Nonequilibrium Biomolecular Dynamics: Exemplified on Amyloid Conformational Dynamics Driven by an Oscillating Electric Field”. Journal of Chemical Theory and Computation 14.7 (July 2018), pp. 3579–3594.

[48] Laurens van der Maaten and Geoffrey Hinton. “Visualizing Data using t-SNE”. Journal of Machine Learning Research 9 (2008), pp. 2579–2605.

[49] A. Bravais. Analyse mathématique sur les probabilités des erreurs de situation d’un point. Impr. Royale, 1844.

[50] Y. Yamamoto, R. Mizuno, T. Nishimura, et al. “Cloning and expression of myelin-associated oligodendrocytic basic protein. A novel basic protein constituting the central nervous system myelin”. The Journal of Biological Chemistry 269.50 (Dec. 1994), pp. 31725–31730.

[51] A Holz and M. E Schwab. “Developmental expression of the myelin gene MOBP in the rat nervous system”. Journal of Neurocytology 26.7 (July 1997), pp. 467–477.

[52] Jacqueline R. Thompson, Erik D. Nelson, Madhavi Tippani, et al. “An integrated single-nucleus and spatial transcriptomics atlas reveals the molecular landscape of the human hippocampus”. Nature Neuroscience 28.9 (Sept. 2025), pp. 1990–2004.

[53] Blue B. Lake, Song Chen, Brandon C. Sos, et al. “Integrative single-cell analysis of transcriptional and epigenetic states in the human adult brain”. Nature Biotechnology 36.1 (Jan. 2018), pp. 70–80.

[54] Matthias Koenning, Stacey Jackson, Curtis M. Hay, et al. “Myelin Gene Regulatory Factor Is Required for Maintenance of Myelin and Mature Oligodendrocyte Identity in the Adult CNS”. The Journal of Neuroscience 32.36 (Sept. 2012), pp. 12528–12542.

[55] M Poncz, S Surrey, P LaRocco, et al. “Cloning and characterization of platelet factor 4 cDNA derived from a human erythroleukemic cell line”. Blood 69.1 (Jan. 1987), pp. 219–223.

[56] Rh Wenger, An Wicki, A Walz, et al. “Cloning of cDNA coding for connective tissue activating peptide III from a human platelet-derived lambda gt11 expression library”. Blood 73.6 (May 1989), pp. 1498–1503.

[57] Aaron Grattafiori, Abhimanyu Dubey, Abhinav Jauhri, et al. The Llama 3 Herd of Models. Version Number: 3. 2024.

[58] Mahmud Omar, Reem Agbareia, Benjamin S Glicksberg, et al. “Benchmarking the Confidence of Large Language Models in Answering Clinical Questions: Cross-Sectional Evaluation Study”. JMIR Medical Informatics 13 (May 2025), e66917–e66917.

[59] Daniel Yang, Yao-Hung Hubert Tsai, and Makoto Yamada. On Verbalized Confidence Scores for LLMs. 2024.

